# α-catenin phosphorylation is elevated during mitosis to resist apical rounding and epithelial barrier leak

**DOI:** 10.1101/2024.09.06.611639

**Authors:** Phuong M. Le, Jeanne M. Quinn, Annette S. Flozak, Adam WT Steffeck, Che-Fan Huang, Cara J. Gottardi

**Author notes:** Co-first authorship (Phuong M. Le, Jeanne M. Quinn). Corresponding Author: Cara J. Gottardi Northwestern University Feinberg School of Medicine 303 Superior St., Simpson-Querrey Institute, 5-525 Chicago, IL 60611.

## Abstract

Epithelial cell cohesion and barrier function critically depend on α-catenin, an actin-binding protein and essential constituent of cadherin-catenin-based adherens junctions. α-catenin undergoes actomyosin force-dependent unfolding of both actin-binding and middle domains to strongly engage actin filaments and its various effectors, where this mechanosensitivity is critical for adherens junction function. We previously showed that α-catenin is highly phosphorylated in an unstructured region that links mechanosensitive middle- and actin-binding domains (known as the P-linker region), but the cellular processes that promote α-catenin phosphorylation have remained elusive. Here, we leverage a previously published phospho-proteomic data set to show that the α-catenin P-linker region is maximally phosphorylated during mitosis. By reconstituting α-catenin Crispr KO MDCK with wild-type, phospho-mutant and mimic forms of α-catenin, we show that full phosphorylation restrains mitotic cell rounding in the apical direction, strengthening interactions between dividing and non-dividing neighbors to limit epithelial barrier leak. Since major scaffold components of adherens junctions, tight junctions and desmosomes are also differentially phosphorylated during mitosis, we reason that epithelial cell division may be a tractable system to understand how junction complexes are coordinately regulated to sustain barrier function under tension-generating morphogenetic processes.

## INTRODUCTION

Simple epithelia comprise a single layer of cells organized into sheets, where they form versatile barriers that compartmentalize tissue organization and functions across organ systems. A key feature that allows individual epithelial cells to form such barriers are intercellular adhesive junctions, which coordinate the coupling of cytoskeletal networks across cells (via adherens junctions and desmosomes), and passage of small molecule constituents between apical and basolateral compartments (via tight junctions) (Angulo-Urarte et al., 2020; Broussard et al., 2020; Citi, 2019; Yap et al., 2018). Since organismal development initiates from the expansion and rearrangement of cells within epithelial sheets, and environmental insults can activate epithelial repair programs, a key question in the field is how cell-cell junction complexes are regulated to allow for dynamic cell-cell behaviors while maintaining overall barrier integrity (Higashi et al., 2024). Indeed, a major challenge in understanding cell-cell adhesion regulation is identifying a well-defined morphogenetic process where complementary proteomic data are also available.

Epithelial cell division is emerging as an ideal system to understand cell-cell junction regulation, as cells dividing in an epithelium undergo defined membrane shape changes, such as apically directed rounding and retraction from the basement membrane to accommodate the mitotic spindle (McKinley et al., 2018), to partitioning cytoplasm via cytokinesis (Derksen and van de Ven, 2020; van de Ven et al., 2016; Wolf et al., 2006) and resolving the midbody through an apical junction abscission mechanism (Bai et al., 2020; Daniel et al., 2018; Herszterg et al., 2014; Higashi et al., 2016; Morais-de-Sa and Sunkel, 2013a; Morais-de-Sa and Sunkel, 2013b). In vertebrate systems, this entire sequence occurs with continuous connection of adherens and tight junction constituents to the actomyosin contractile ring during cytokinesis and full maintenance of the transepithelial barrier (Higashi et al., 2016), suggesting epithelial junctions can withstand mitotic forces.

Recent studies suggest that adherens junctions (AJs), particularly the cadherin-catenin adhesion complex and its essential actin-binding component α-catenin (α-cat), may be a central mechanosensitive mediator of epithelial cell division. In cleaving Xenopus embryos, E-cadherin and β-catenin proteins showed reduced mobility at the cytokinetic furrow relative to non-dividing membrane interface, along with enhanced recruitment of the vinculin, a homologue and mechanosensitive binding partner of α-cat (Higashi et al., 2016). Related work in dividing MDCK epithelial monolayers revealed that as a mitotic cell rounds up and away from its neighbors, it generates increased tension on an adjacent cell’s junctions, favoring vinculin recruitment (Monster et al., 2021). This asymmetric recruitment of vinculin to AJs in neighboring, rather than dividing cells, contributes to epithelial barrier integrity, as MDCK cells reconstituted with an α-cat mutant that cannot recruit vinculin showed clear gaps and barrier leak when present in neighboring, rather than mitotic cells. Together, these data suggest that the cadherin-catenin complex is mechanically altered during mitosis to promote effector (e.g., vinculin) recruitment to preserve epithelial barrier integrity. Whether adherens junction regulation during cell division largely relies on force-dependent unfolding of α-cat, independent of other modes of regulation, is not known. In the study that follows, we show that α-cat phosphorylation is upregulated during mitosis and contributes to epithelial barrier function in MDCK cells. Along with previously published phospho-proteomic data sets showing that major scaffold components of adherens junctions, tight junctions and desmosomes are differentially phosphorylated during mitosis (Dephoure et al., 2008), we reason that epithelial cell division may be a tractable system to understand how adhesive junction complexes are regulated.

## RESULTS

### α-cat phosphorylation is increased during mitosis

Quantitative phosphoproteome profiling of various cell and tissue systems confirmed evidence by our group that α-catenin is reproducibly phosphorylated at multiple sites in an unstructured region that links mechanosensitive middle- and actin-binding domains (Ballif et al., 2004; Beausoleil et al., 2004; Dephoure et al., 2008; Escobar et al., 2015; Huttlin et al., 2010; Olsen et al., 2006; Zhai et al., 2008). While in vitro kinase assays using purified recombinant α-cat as substrate established a Casein Kinase 2 (CK2)-Casein Kinase 1 (CK1) dual-kinase mechanism ((Escobar et al., 2015); Fig. 1A), upstream signals and processes that regulate α-cat phosphorylation remained elusive. Curiously, stable isotope labeling of HeLa cells arrested in the G_1_ or Mitotic phases of the cell cycle suggest α-cat phosphorylation as quantitatively increased during mitosis (Dephoure et al., 2008) (Fig. 1A), but reproducibility of this regulation and its role in epithelial cell division are lacking. We used commercially available antibodies that recognize distinct α-cat phospho-sites to immunoblot lysates prepared from HeLa cells synchronized in G1/S or G2/M phases of the cell cycle (Fig. 1B). We found that α-cat phosphorylation at S641 is not obviously enhanced by mitosis, whereas α-cat phosphorylated at S652 or S655/T658 clearly increases during mitosis (Fig. 1B-C). These data suggest mitosis does not impact α-cat phospho-priming at the most abundant site (S641), but rather increases phosphorylation at previously defined CK1 sites (pS652, pS655/T658), which are sequentially related (Escobar et al., 2015). Since this phospho (P)-domain resides in a region that links middle-(M) and actin-binding domains (ABD), we refer to this as the P-linker region (Escobar et al., 2015).

**Fig. 1:**
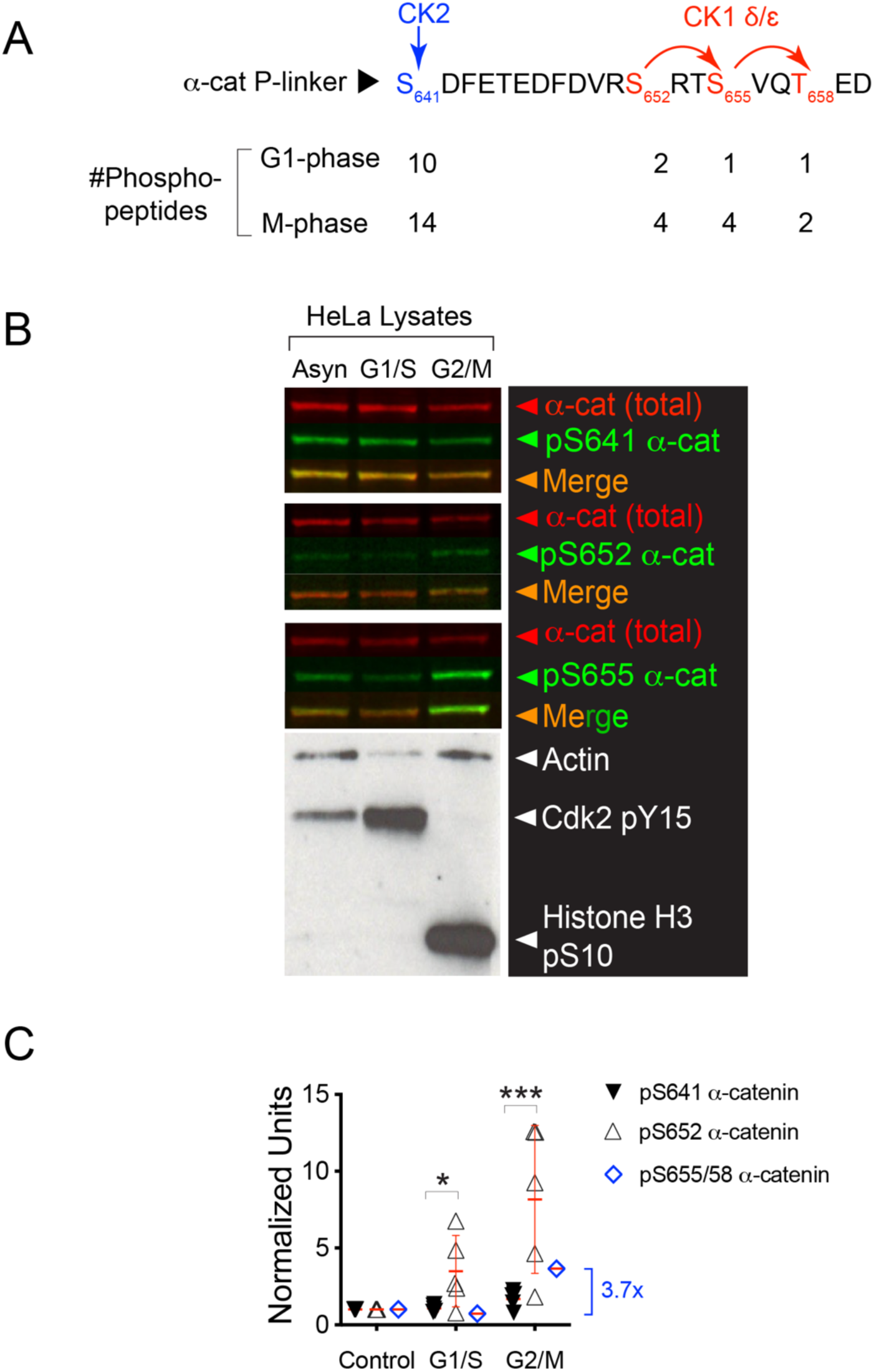
α-cat phosphorylation is increased during mitosis in synchronized HeLa cell lysates. (A) Phospho-peptide detection during G1- and mitotic (M) phases of the cell cycle by stable isotope labeling phospho-peptide enrichment mass spectrometry (Dephoure et al., 2008). Schematic shows previously defined in vitro dual-kinase mechanism, where CK2 phosphorylation at S641 primes α-cat for subsequent and sequential phosphorylations by CK1 at S652, S655 and T658 (Escobar et al., 2015). **(B)** Immunoblot validation of α-cat phosphorylations at S641 (Signalway), S652 and S655/T658 (Cell Signaling) in asynchronized and synchronized G1/S and G2/M HeLa cell lysates. Actin, pY15 Cdk2 and pS10-Histone 3 are used as loading controls to validate cell cycle phases. **(C)** Quantification of α-cat phospho-site detection from multiple immunoblots (pS641 and pS652). Significance by 2-way ANOVA, *** (p = 0.0003) and * (p = 0.04). Single blot shown for pS655/T658 reveals 3.7-fold increase in phosphorylation relative to the G1/S condition.

### Phospho-mimic α-cat restrains mitotic apical rounding

To address consequences of α-cat P-linker phosphorylation for mitosis, we restored α-cat CRISPR-knock-out Madin Darby Canine Kidney (MDCK) cells (Quinn et al., 2024) with GFP-tagged α-catenin proteins, where previously mapped phosphorylations were blocked or charge-mimicked by amino acid substitution (α-cat 4A and 4E mutants, respectively) (Fig. 2). Newly confluent MDCK monolayers grown on glass coverslips (48hrs) were fixed, DNA-stained and imaged to quantify epithelial cell shape changes during established phases of cell division (metaphase, anaphase and telophase), which we reasoned might be altered by α-cat P-linker modification state. By tracing mitotic cell perimeters, we found that α-cat phospho-mimic (4E) cells appear significantly larger than α-cat phospho-mutant (4A) cells (Fig. 3A; Fig. S1). This apparent difference in mitotic cell area is not due to intrinsic differences in cell size (Fig. S2). Instead, we found that α-cat phospho-mimic (4E) -restored MDCK cells show less apical rounding than wild-type or phospho-mutant (4A) -expressing cells (Fig. 3B-C, orthogonal x-z views). Indeed, the apical surface of newly confluent α-cat phospho-mimic (4E) epithelial monolayers appeared taut and generally flatter than wild-type or phospho-mutant (4A) -expressing cells; conversely, the cortex of mitotic α-cat 4A cells appeared slack, following the contours of condensed chromosomes and nuclei (Fig 3B, arrows; 3D-Video 1). These data suggest that full phosphorylation of α-cat’s P-linker region constrains mitotic rounding within the epithelial monolayer and generally promotes epithelial maturation.

**Fig. 2:**
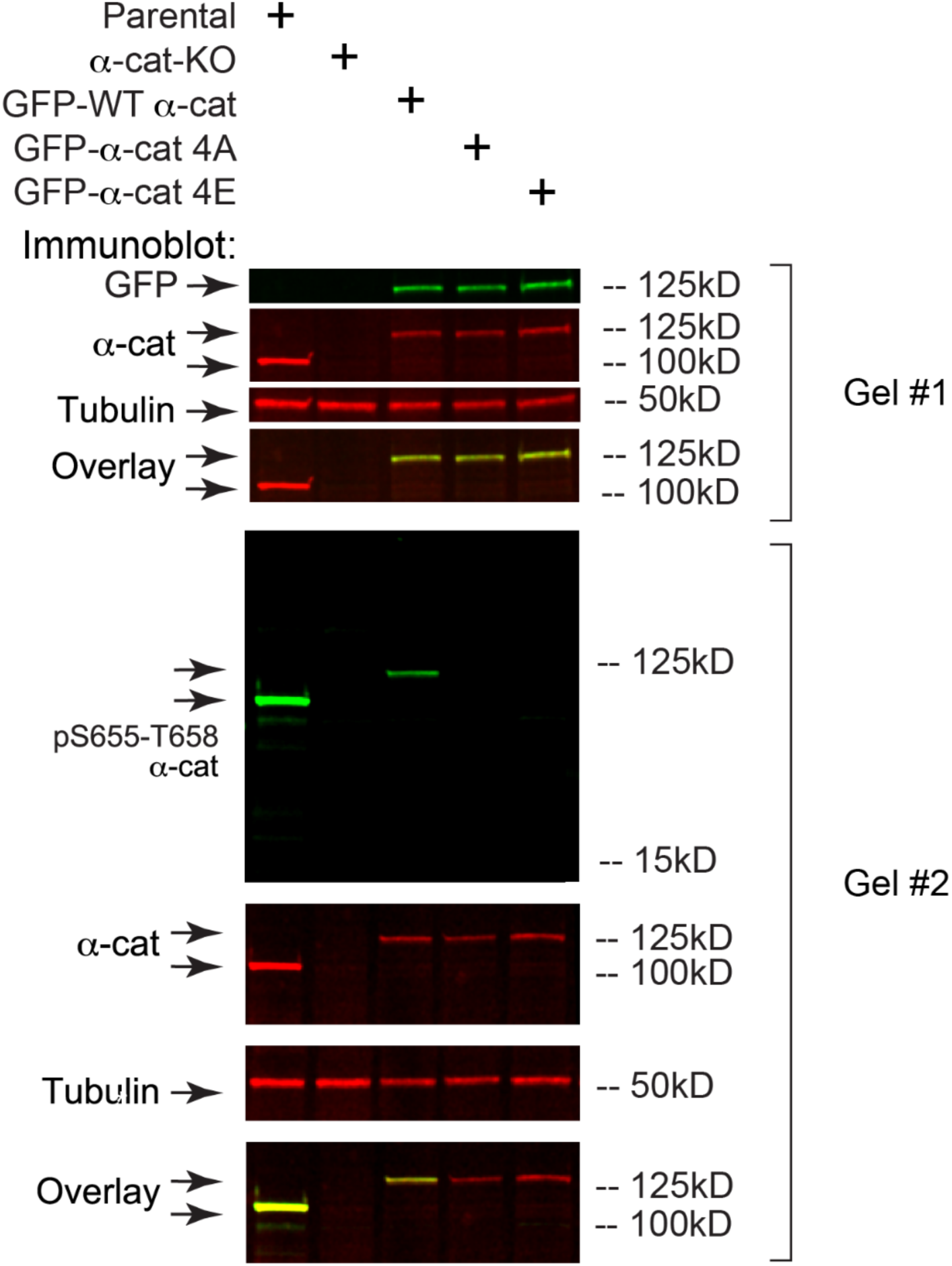
GFP-α-cat mutants express α-cat similarly in α-cat KO MDCK cells. Immunoblot validation of α-cat and GFP-α-cat in MDCK cell line parentals, α-cat-KO, α-cat-KO^GFP-α-cat^ ^WT^, α-cat-KO^GFP-α-cat^ ^4A^, and α-cat-KO^GFP-α-cat^ ^4E^. Tubulin is used as a loading control to validate loading protein amount (Gel #1). Antibodies to α-cat phosphorylated at S655/T658 do not recognize α-cat 4A or 4E mutant constructs, as expected (Gel #2).

**Fig. 3:**
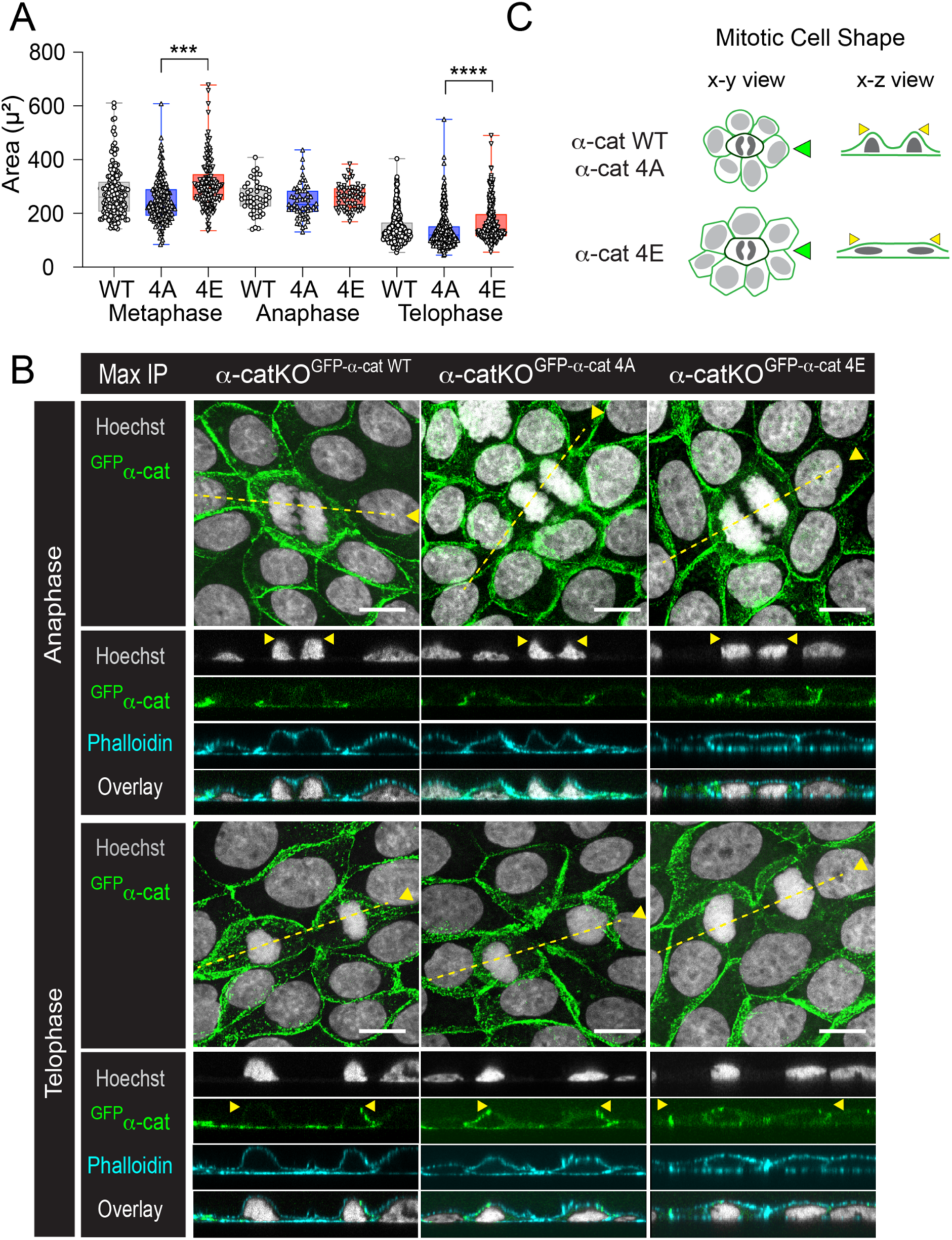
Phospho-mimic α-cat restrains mitotic rounding compared with wild-type and phospho-mutant α-cat. **(A)** Quantification of cell area (microns^2^) of α-cat cell lines during mitotic phases. Data presented as mean ±SD with significance by ANOVA, **** (p < 0.0001) and ***(p = 0.0003). Image captured using a 20x objective Nikon Ti2a microscope. **(B)** Confocal images (z-stack maximum intensity projection, MIP) taken on AXR Nikon microscope of MDCK monolayer fixed and stained for DNA (Hoechst, gray), F-actin (Phalloidin, cyan) or α-cat (native GFP fluorescence, green). Overlay image with complementary orthogonal x-z view along mitotic cell (dotted yellow line) shows apical extension of nucleus during mitosis. Scale bar = 10μm. **(C)** Schematic of a mitotic cell (dark green) on neighboring cells’ membrane (light green) during mitotic phases between α-cat mutants. Mitotic α-cat-KO^GFP-α-cat^ ^4A^ cells measured the smallest cell area suggesting rounding in the z-direction. Green arrows indicate x-z side view; yellow arrowheads rationalize area quantification differences in A and B.

### Phospho-mimic α-cat reduces barrier leak during telophase

Mitosis generally relies on actomyosin contractility-dependent rounding to accommodate spindle formation for chromosome segregation and cytokinesis into genetically identical daughters (Taubenberger et al., 2020). Epithelia need to execute these steps in a manner that preserves interactions with neighbors to maintain the barrier, a key function of epithelia across tissue types (Higashi et al., 2016). We wondered, therefore, if α-cat phosphorylation in the P-linker might limit intercellular junction leak, particularly between mitotic cells and their non-dividing neighbors. We used an established assay to visualize small or transient intercellular leaks, which seeds epithelial cells on a biotinylated collagen matrix at confluent density, and reveals monolayer breach via fluorescent conjugated-streptavidin (Dubrovskyi et al., 2013; Monster et al., 2021) (Fig. S3A Schematic). We observed many leaks in wild-type or phospho-mutant (4A) -restored MDCK cells, particularly during telophase where actomyosin forces may be peaking. Very few breaks were detected in α-cat phospho-mimic (4E) -restored MDCK cells (Fig. 4). Qualitatively, leak size (area) was greater for α-cat wild-type and phospho-mutant (4A) than α-cat phospho-mimic (4E)-restored MDCK cells (Fig. 4B-D). These data suggest that full phosphorylation of α-cat’s P-linker region promotes epithelial barrier integrity during mitosis by strengthening interactions between dividing and non-dividing neighbors. α-cat phosphorylation also appears to play a more general role in epithelial barrier integrity (Quinn et al., In preparation).

### Phosphorylated α-cat localizes to the apical most portion of epithelial cell junctions

HeLa cell phospho-proteomic and α-cat phospho-antibody immunoblot data reveal that the α-cat P-linker region is maximally phosphorylated during mitosis (Fig. 1). Since HeLa cells synchronized in mitosis are released from tissue culture plates after rounding (i.e., double-thymidine block, post-nocodazole mitotic “release” method; (Dephoure et al., 2008), it is likely that the increase in α-cat phosphorylation occurs within the mitotic cell itself, rather than via neighboring cells (i.e., a mitotic cell autonomous versus non-autonomous mechanism). We wondered, therefore, whether we could determine subcellular localizations of phospho-specific forms of α-cat in dividing MDCK cells using available antibodies (Fig. 1). We chose to assess phospho-α-cat localization in MDCK, rather than HeLa cells, since the latter are derived from a poorly differentiated adenocarcinoma and not strongly self-adherent (Doyle et al., 1995), although adherens-like structures have been described (Deng et al., 2008; Izawa et al., 2002; Pestonjamasp et al., 1997). Interestingly, antibodies that recognize terminal phosphorylations in the α-cat CK1 sequence, pS655 and T658 (Escobar et al., 2015), decorate cell-cell junctions of both dividing and non-dividing MDCK cells (Fig. 5). Confocal imaging shows that antibodies to phospho-α-cat largely overlap with an antibody that recognizes total α-cat (Fig. 5A, top row). Curiously, optical sections in the x-z direction show that the phospho-α-cat signal appears to specifically decorate apical junctions (Fig. 5A, magenta/green arrows; see also inset (i)). Since antibodies to α-cat pS655/T658 also showed an extra-junctional punctate staining pattern (asterisks), possibly elevated in mitotic cells, we used a proximity ligation assay (PLA) to validate the localization of phospho-α-cat in MDCK cells (Fig. 5B-C). This method allowed us to use PLA as a “coincidence-detection system” for total and phospho-α-cat, amplifying the cellular localization of phospho-α-cat and reducing impact of antibody cross-reactivity with other possible pS/T epitopes (although we note that the pS655/T568 antibody does not detect cross-reactive bands across a wide molecular weight range by immunoblot analysis, Fig. 2). While the amplified proximity signal of total α-cat/phospho-α-cat is sparse compared to indirect immunofluorescence methods, this method appears to selectively detect phospho-α-cat at cell-cell junctions (Fig. 5B-C). Curiously, optical sections in the x-z direction show that the total α-cat/phospho-α-cat proximity signal is at the apical-most portion of cell-cell junctions (Fig. 5D). Proximity detection using antibodies to pS641 and pS652 α-cat show similar apical bias (Fig. 5D, lower panels). Note that the MDCK monolayer in Figure 5A was grown on glass, in contrast to filter-grown cells in 5D, which may contribute to the greater apical enrichment of phospho-α-cat in latter images. Surprisingly, we saw no obvious increase in proximity-amplified total α-cat/phospho-α-cat signal in dividing versus non-dividing MDCK cells (Fig. 5B&D, yellow arrows). These data are in contrast with the increased abundance of phospho-α-cat detected in mitotic HeLa cells (Fig. 1)(Dephoure et al., 2008), and suggest that the extent of α-cat phosphorylation may be more related to a feature common to mitotic HeLa cells and MDCK cell monolayers.

**Fig. 4:**
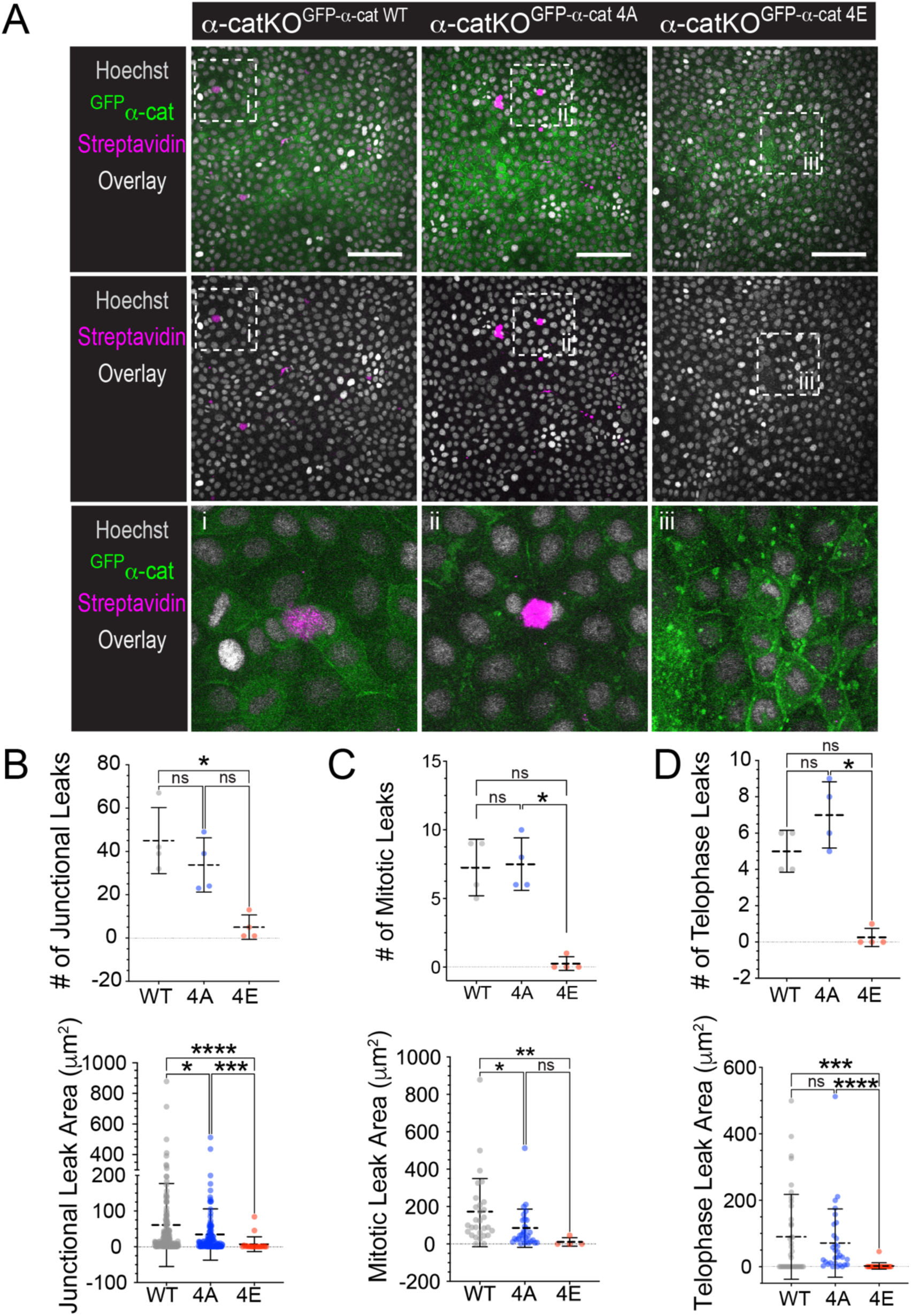
Phospho-mimic α-cat reduces barrier leak during mitotic rounding compared with α-cat-WT and α-cat-4A. **(A)** Confocal image (z-stack maximum intensity projection of basal region) taken on AXR Nikon microscope of MDCK permeability assay fixed and stained for DNA (Hoechst, gray), Streptavidin binding to biotinylated collagen (magenta) and α-cat (native GFP, green). Overlay image shows basal biotin-streptavidin interactions in α-cat-KO^GFP-α-cat^ ^WT^ and α-cat-KO^GFP-α-cat^ ^4A^ during telophase (white box inset, i, ii and iii). Scale bar = 100μm. **(B)** Quantification of leak area and total junctional leak between α-cat mutants. Data presented as mean ±SD with significance by ANOVA, ****(p < 0.0001), *** (p = 0.0003), and * (p = 0.0122). **(C)** Quantification of leak area and total mitotic leak between α-cat mutants. Data presented as mean ±SD with significance by ANOVA, **(p = 0.0233) and * (p = 0.0028). (**D)** Quantification of leak area and total telophase leak between α-cat mutants. Data presented as mean ±SD with significance by ANOVA, ****(p < 0.0001), *** (p = 0.0003), and * (p = 0.0122).

**Fig. 5:**
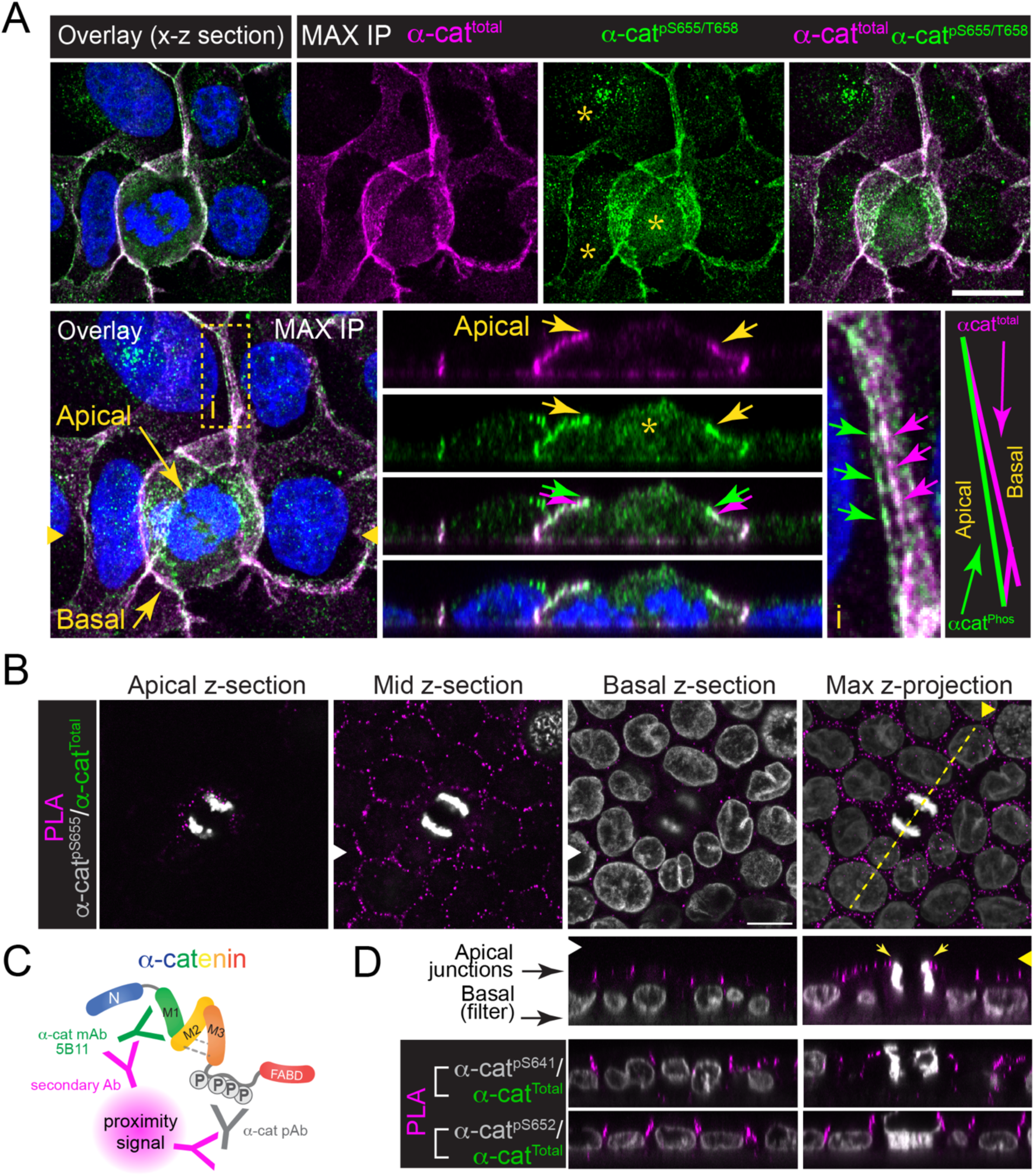
Phospho-α-cat localizes to the apical most portion of epithelial cell junctions. (A) Confocal image (z-stack maximum intensity projection, MIP) of MDCK monolayer (glass coverslip grown) fixed and immuno-stained with antibodies to α-cat (magenta) and α-cat phosphorylated at S655/T658 (green). DNA stained with Hoechst (blue). Overlay image with complementary orthogonal x-z views shows pS655/T658 apical junction enrichment (yellow arrows) in both mitotic cell and adjacent cell junctions (yellow box inset, i). Asterisk (*) shows punctate cytoplasmic staining with pS655/T658 antibody that is likely non-specific. **(B)** Confocal x-y sections of α-cat pS655/T658 co-incidence detection (magenta spots) using proximity ligation assay (PLA) on filter-grown MDCK cells (10 days). DNA in gray. **(C)** Schematic of proximity ligation assay (PLA) using two antibodies to α-cat. **(D)** Orthogonal x-z sections of α- cat/pS655/658, α-cat/pS641 and α-cat/pS652 co-incidence detection (magenta spots). Scale bar = 10μm.

## DISCUSSION

While the cadherin-catenin complex is long known to be required for adherens junction (AJ) organization and epithelial barrier homeostasis (Gumbiner et al., 1988), we know comparatively less about how and under what conditions the cadherin-catenin complex is regulated. A major paradigm shift in thinking about adherens junction regulation is that the cadherin-catenin complex is mechanosensitive, particularly via its essential actin-binding component, α-catenin (Angulo-Urarte et al., 2020). Indeed α-cat’s actin-binding and middle-domains undergo force-dependent unfolding to engage F-actin or various actin-binding effectors (e.g., vinculin), respectively (Barrick et al., 2018; Buckley et al., 2014; Kim et al., 2015; Twiss et al., 2012; Wang et al., 2022; Yao et al., 2014; Yonemura et al., 2010). This allows actomyosin-force dependent strengthening of α-cat binding to actin via direct and indirect mechanisms. Curiously, α-cat is not only regulated by force; α-cat is highly phosphorylated in an unstructured region that links mechanosensitive middle- and actin-binding domains (known as the P-linker region) (Escobar et al., 2015). While previous *in vitro* kinase assays revealed an elaborate dual-kinase mechanism, where phosphorylation at S641 by CK2 effectively primes α-cat for further sequential phosphorylation at S652, S655 and T658 by CK1 (Escobar et al., 2015), the cellular processes and upstream kinase/phosphatase signals that promote α-cat phosphorylation *in vivo* remained unknown.

Here, we leveraged previously published high throughput phospho-proteomics data (Dephoure et al., 2008) to show that phosphorylation of α-cat’s P-linker region is elevated during mitosis, particularly at previously characterized CK1 sites (Escobar et al., 2015), using HeLa cell synchronized lysates and validated phospho-specific antibodies to α-cat (Cell Signaling). Since HeLa cells are cancer-derived and not typically used for studying cell-cell adhesion, we sought to validate the role of α-cat phosphorylation during mitosis in MDCK cells, a longstanding model to study epithelial junctions (Dukes et al., 2011). By reconstituting α-cat Crispr KO MDCK with wild-type, phospho-mutant (4A) or phospho-mimic (4E) forms of GFP-α-cat, we show that amino acid charge substitution of α-cat’s P-linker, which aims to mimic the full and persistent phosphorylation of α-cat, constrains mitotic division within the plane of an MDCK epithelial monolayer, limiting intercellular breaks that form between dividing and non-dividing cells. We also observed that wild-type and α-cat phospho-mutant-restored nascent MDCK monolayers appear generally leakier and less mature than the α-cat phospho-mimic line, with the former showing a more “fried-egg” morphology with compliant apical membranes overlying the nucleus. These data suggest that full phosphorylation of the α-cat P-linker region may also be generally required for epithelial monolayer shape-transitions that lead to a mature barrier (Cammarota et al., 2024). Overall, while these data are in line with our previous work showing α-cat phosphorylation contributes to epithelial monolayer adhesive strength and cell-cell coordination during collective migration (using an α-cat shRNA MDCK knock-down/GFP-α-cat reconstitution system (Escobar et al., 2015), they advance an important new concept—α-cat phosphorylation is not simply constitutive, but can increase during mitotic morphogenesis to maintain epithelial barrier function under strain.

We do not yet understand how mitotic signaling causes the upregulation of α-cat phosphorylation at CK1 sites. We previously discovered that the CK1 sites in α-cat are less accessible to in-solution phosphorylation by CK1 in full length α-cat compared with a fragment comprising only the C-terminal half of α-cat ((Escobar et al., 2015), Fig. 2G-H of that paper). This raises the possibility that α-cat binding to actin or increased actomyosin contractility associated with mitosis might favor α-cat P-linker unfolding and kinase accessibility. However, we cannot exclude the possibility that mitosis upregulates other kinases or inhibits phosphatases that target α-cat at S652, S655 and T658.

We also do not fully understand how α-cat phosphorylation reinforces epithelial barriers during cell division. Recent studies implicate vinculin, an α-cat homologue and mechanosensitive binding partner as a key adherens junction reinforcer during cell division (Higashi et al., 2016; Monster et al., 2021). In Xenopus, vinculin becomes enriched along the cytokinetic furrow, coincident with a reduction in cadherin/catenin mobility (Higashi et al., 2016). Since loss of vinculin or its coupling to actin enhances the rate of furrow ingression and tight junction leak (Higashi et al., 2016)(van den Goor et al., BioRxiv 2023), it appears that the speed of mitosis/cytokinesis must be carefully controlled by the cadherin-catenin complex (Goldbach et al., 2010; Padmanabhan et al., 2017) to ensure epithelial barrier maintenance during cell division. Of interest, evidence from MDCK cells suggests that mitotic force-dependent α-cat-unfolding and recruitment of vinculin appears to be asymmetric, requiring reinforcement of adherens junctions by vinculin in cells surrounding, rather than within the mitotic cell (Monster et al., 2021). Given these data, it is attractive to speculate that α-cat phosphorylation-dependent epithelial barrier reinforcement during cell division may be due, at least in part, to enhanced vinculin recruitment. However, since we previously found that an α-cat phospho-mimic mutant incapable of binding vinculin could not reverse cell-cell cohesive behaviors enhanced by phosphorylation (Escobar et al., 2015), α-cat phosphorylation likely impacts α-cat structure-function more broadly, and beyond simply recruiting vinculin (Quinn et al., manuscript in progress).

Evidence that a mitotic cell rounding against its neighbor can lead to adherens junction asymmetry (Monster et al., 2021) inspired us to look closely at where phospho-α-cat is localized in dividing MDCK cells. While phosphorylation of α-cat’s P-linker region is clearly elevated in mitotic HeLa cell lysates, we saw no clear increase in phospho-α-cat detection along the dividing/non-dividing MDCK adherens junction. Instead, we found that phospho-α-cat appears localized to adherens junctions more generally, and notably the apical most region of adherens junctions known as the zonula adherens (Mangeol et al., 2024; Mooseker et al., 1983). Similar immunofluorescence analysis in HeLa cells was not possible, possibly because this cancer-derived cell line is known to make only weak spot-like adherens junctions (Deng et al., 2008; Izawa et al., 2002; Pestonjamasp et al., 1997) (not shown). We speculate, therefore, that full phosphorylation of the α-cat P-linker region may depend on a property common to mitosis and zonula adherens junctions, such as a reliance on actomyosin-based contractility (Murrell et al., 2015; Nyga et al., 2023; Sorce et al., 2015; Yap et al., 2018).

In summary, these data suggest that full phosphorylation of α-cat’s P-linker region promotes epithelial barrier integrity during mitosis by strengthening interactions between dividing and non-dividing neighbors. Along with previously published phospho-proteomic data sets showing that major scaffold components of adherens junctions, tight junctions and desmosomes are differentially phosphorylated during mitosis ((Dephoure et al., 2008; Olsen et al., 2010); Table 1), we reason that epithelial cell division may be a tractable system to understand how junction complexes are coordinately regulated.

**TABLE 1.**

| GENE/Protein | HeLa cells |  |
| --- | --- | --- |
|  | S-phase | Mitosis |
|  | # of phospho-sites |  |
| **Adherens Junctions** |  |  |
| CTNNA1/α-catenin | 5 | 5* |
| CTNNB1/β-catenin | 0 | 2 |
| CTNND1/p120<sup>ctn</sup> | 5 | 9 |
| ARVCF/δ-catenin | 1 | 2 |
| PKP4/ Plakophilin4/p0071 | 1 | 11 |
| MLLT4 (AFDN)/Afadin | 6 | 10 |
| **Tight Junction** |  |  |
| TJP1/ZO-1 | 6 | 22 |
| TJP2/ZO-2 | 17 | 25 |
| TJP3/ZO-3 | 3 | 4 |
| PARD3/Par3 | 4 | 9 |
| CGN/Cingulin | 5 | 5* |
| CLDN12/Claudin 12 | 0 | 2 |
| F11R/Jam1 | 6 | 2 |
| MARVELD2/Tricellulin | 2 | 0 |
| NF2/Merlin | 1 | 0 |
| JUB/Ajuba | 1 | 0 |
| **Desmosome** |  |  |
| DSG2/Desmoglein | 1 | 2 |
| DSP/Desmoplakin | 29 | 19 |
| PKP2/Plakophilin2 | 6 | 3 |
| PKP3/ Plakophilin3 | 5 | 4 |
| *Some phospho-sites are different between S- and M-phase; others are increased in M-phase |  |  |
| Color shades within junction systems indicates more or less phosphorylation |  |  |
| Data selected from Table S1 of Dephoure et al. [Gygi], A quantitative atlas of mitotic phosphorylation. PNAS 105; 31, 2008. |  |  |

## ACKNOWLEDGEMENTS

This work relied on the following Northwestern University services and core facilities: Flow Cytometry (NCI CA060553); Center for Advanced Microscopy (NCI CCSG P30 CA060553) and Skin Biology and Diseases Resource-based Center (P30AR075049) from the NIAMS. We thank David Kirchenbeuchler (Northwestern Imaging Facility) for advice. This work would not able been possible without longstanding use of the HeLa cell line. We gratefully acknowledge Henrietta Lacks, and the Lacks family, for their contribution to biomedical research.

## FUNDING

JMQ is supported by T32 HL076139 and F30EY036267; CJG by NIH GM129312 and HL163611. All authors declare no competing financial interests.

### Author contributions

PML, JMQ, ASF, AWTS and CFH conducted experiments and analyzed results; CJG designed and supervised the study. PML, JMQ and CJG wrote the manuscript. CJG provided funding for the project.

## METHODS

### Plasmid constructs

N-terminally GFP-tagged αE-catenins were synthesized by VectorBuilder using a dimerization-disrupted mEGFP (A206K) in third-generation lentiviral vectors with components pLV[Exp]-CMV>mEGFP-αE-catenin EF1A(core)>Puro. Lentivirus packaging (psPAX2, #12260) and envelope (pMD2.G, #12259) plasmids were purchased from Addgene. Previously established α-cat phospho-sites S641, S652, S655 and T658 (Escobar et al., 2015) were changed to alanine (α-cat 4A mutant, which prevents phosphorylation) or glutamate (α-cat 4E mutant, which aims to mimic the phosphate charge).

### Cell culture and stable cell line selection

MDCK II cells were maintained in Dulbecco’s Modified Eagle’s Medium (DMEM, Corning), containing 10% fetal bovine serum (FBS, R&D Systems), 100 U/ml penicillin and 100 μg/ml streptomycin (Corning). α-cat/*Ctnna1* knockout MDCK clone 2.2 was generated using CRISPR-Cas9 system as described in Quinn et al (Quinn et al., 2024). For lentivirus production, 293T cells (GeneHunter) were transfected with 8μg expression vector (Vector Builder), 6μg psPAX2, and 2μg pMD2.G using TransIT (Mirus). Viral supernatant was collected 48 and 72h after transfection, passed through a 0.45μm filter and supplemented with 1μL/mL polybrene (Sigma). To generate stable GFP-α-cat lines, MDCK α-cat KO cells were transduced for 6hr at 37°C on 10cm plates with 2mL prepared viral supernatant. Cells were selected in culture media containing 5μg/mL puromycin, then sort-matched for GFP using a FACS Melody 3-laser sorter (BD).

### Antibodies

The following primary antibodies were used: polyclonal rabbit anti-α-cat (C3236, Cell Signaling), hybridoma mouse anti-α-catenin (5B11, (Daugherty et al., 2014)), polyclonal rabbit anti-GFP (A11122, Invitrogen) and Phalloidin-488 or -568 (A12379, Invitrogen). Secondary antibodies for Western blotting included HRP-conjugated goat anti-mouse and -rabbit antibodies (Bio-Rad), or fluorescently labeled donkey anti-mouse and -rabbit antibodies (680RD or 800RD, LiCor Biosciences). Secondary antibodies for immunofluorescence included IgG Alexa Fluor 488 or 568-conjugated goat anti-mouse or -rabbit antibodies (Invitrogen).

### Immunofluorescence and Imaging

Cells were grown on glass coverslips, fixed in 4% paraformaldehyde (Electron Microscopy Services, Hatfield, PA) for 15’, quenched with glycine, permeabilized with 0.3% Triton X-100 (Sigma), and blocked with normal goat serum (Sigma). Primary and secondary antibody incubations were performed at RT for 1h, interspaced by multiple washes in PBS, and followed by mounting coverslips in ProLong Gold fixative (Life Technologies). Images of mitotic GFP-α-cat WT, 4A and 4E -expressing MDCK monolayers were captured with a Nikon Ti2 (B) Widefield Microscope (DS-Qi2 Camera, 20x air objective) using NIS Elements software. Confocal z-stack (0.3μm step size) images were captured using a Nikon AXR with GaAsP detectors and equipped with 95B prime Photometrics camera, Plan-Apochromat 40x objective.

### Image Analysis and Quantification

To examine the α-cat phosphorylation on mitotic rounding, cell area was quantified on maximum intensity projection in FIJI. The area was measured by ROI through hand tracing of cell junctions from the GFP-α-catenin signal. To compare barrier function of GFP-α-cat wild-type, phospho-mutant or -mimic restored MDCK cells, junction leak (streptavidin conjugated with Alexa Fluor 568, below) was quantified from maximum intensity projections of the glass/basal surface through cell height in FIJI. Leak area, perimeter and number were measured in FIJI through hand tracing the biotin-streptavidin signal. All quantifications of mean, standard deviation, and significance by ANOVA was conducted through GraphPad Prism.

### Epithelial Permeability Immunofluorescence Assay

Glass-bottom dishes (Falcon) were coated with 1 mg/mL Collagen IV (C5533, Sigma-Aldrich) for 30 minutes at 37°C. Then, dishes were biotinylated with EZ-Link-NHS-LC-Biotin (21336, Thermo Fisher Scientific) at 1.5 mg/mL overnight at 4°C. MDCK α-cat knockout cells expressing GFP-α-cat-WT, GFP-α-cat-4A, or GFP-α-cat-4E were seeded and cultured for 48-hrs to develop a nascent epithelial monolayer. Cells were washed with cold PBS-Ca/Mg++, treated with 25 µg/mL streptavidin conjugated with Alexa Fluor 568 (S11226, Thermo Fisher Scientific) for 30 minutes at 4°C before being rinsed with PBS, fixed and processed for immunostaining.

## Key Resources Table

| REAGENT or RESOURCE | SOURCE | IDENTIFIER |
| --- | --- | --- |
| **Antibodies** |  |  |
| α-catenin 15D9 | Enzo Life Sciences | ALX-804-101 |
| α-catenin pS641 | Signalway | 11330 |
| α-catenin pS652 | Cell Signaling | 13061 |
| α-catenin pS655, 658 | Cell Signaling | 13231 |
| Alexa Fluor 568 Phalloidin | Invitrogen | A12380 |
| Cell Cycle and Apoptosis Western Blot Cocktail | Abcam | ab139417 |
| GFP | Invitrogen | A11122 |
| Streptavidin, Alexa Fluor 568 | ThermoFisher Scientific | S11226 |
| Tubulin | Sigma | T4026 |
| **Chemicals, Peptides, and Recombinant Proteins** |  |  |
| Collagen IV | Sigma-Aldrich | C5533 |
| Dulbecco's Modification of Eagle's Medium (DMEM) with 4.5 g/L glucose, L-glutamine, & sodium pyruvate | Corning | 10-013-CV |
| Dulbecco's Phosphate – Buffered Saline (PBS), 1x, without calcium & magnesium | Corning | 21-031-CV |
| Dulbecco's Phosphate – Buffered Saline (PBS), 1x, with calcium & magnesium | Corning | 21-030-CM |
| EZ-Link-NHS-LC-Biotin | ThermoFisher Scientific | 21336 |
| Fetal Bovine Serum | BioTechne | S11510 |
| HeLa Cell Cycle Lysates | Abcam | ab136811 |
| Penicillin Streptomycin Solution, 50x | Corning | 30-001-CI |
| 0.25% Trypsin, 2.21 mM EDTA, 1X [-] sodium bicarbonate | Corning | 25-053-CI |
| Hoechst 33342 | ThermoFisher Scientific | 62249 |
| ProLong Gold Antifade Mountant | ThermoFisher Scientific | P36934 |
| **Critical Commercial Assays** |  |  |
| **Experimental Models: Organisms/Strains** |  |  |
| Madin-Darby canine kidney II cells | Gumbiner Lab (1996) | Heidelberg strain (Kai Simmons Lab) |
| **Software and Algorithms** |  |  |
| FIJI/ImageJ (version: 2.1.0/1.53c) | Schneider, CA et al., 2012 | https://imagej.nih.gov/ij/ |
| Graphpad Prism |  | https://www.graphpad.com/scientific-software/prism/ |
| Oxford Instruments Imaris (version 10.1) |  | https://imaris.oxinst.com/ |
| **Other** |  |  |
| AXR Confocal | Nikon Instruments | Galvano scanning, GaAsP detectors |
| Axioplan2 Epifluorescence Microscope | Zeiss; 20x objective (Air) | AxioCAM HR Camera with AxioVision 4.8 software |
| Ti2 (B) Widefield Microscope | Nikon Instruments | Wide-field Nikon DS-Qi2 Camera, 20x air objective, NIS Elements |
| Nikon Eclipse TS100 | Nikon Instruments | 10x objective, Apple iPhone15 camera |
| Plasmids |  |  |
| Monomeric GFP-αE-catenin | VectorBuilder | Human α-Ecat sequence/*CTNNA1*; first used in Quinn et al., 2024 |
| Monomeric GFP-αE-catenin P-linker 4A | VectorBuilder | S<sub>641</sub>S<sub>652</sub>S<sub>655</sub>T<sub>658</sub> →<br>A<sub>641</sub>A<sub>652</sub>A<sub>655</sub>A<sub>658</sub> |
| Monomeric GFP-αE-catenin P-linker 4E | VectorBuilder | S<sub>641</sub>S<sub>652</sub>S<sub>655</sub>T<sub>658</sub> →<br>E<sub>641</sub>E<sub>652</sub>E<sub>655</sub>E<sub>658</sub> |

---

**Fig. S1:**
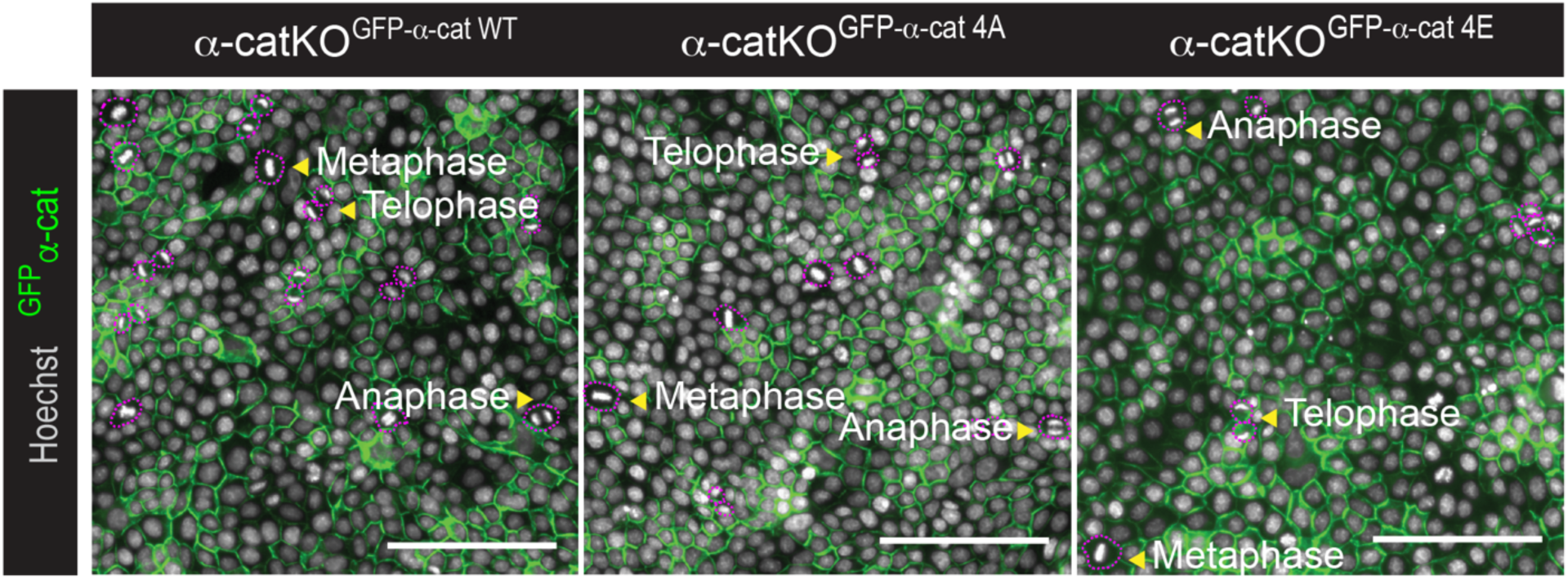
Tracings of dividing MDCK cells. *[Related to Fig. 3]* Nikon Ti2 Widefield images (z-stack maximum intensity projection of basal region) of MDCK fixed and immuno-stained for α-cat (native GFP, green) and Hoechst (gray). Overlay image with cell area hand tracing (dotted magenta) shows criteria for differentiating cell division stage (yellow arrowheads). Scale bar = 100μm.

**Fig. S2:**
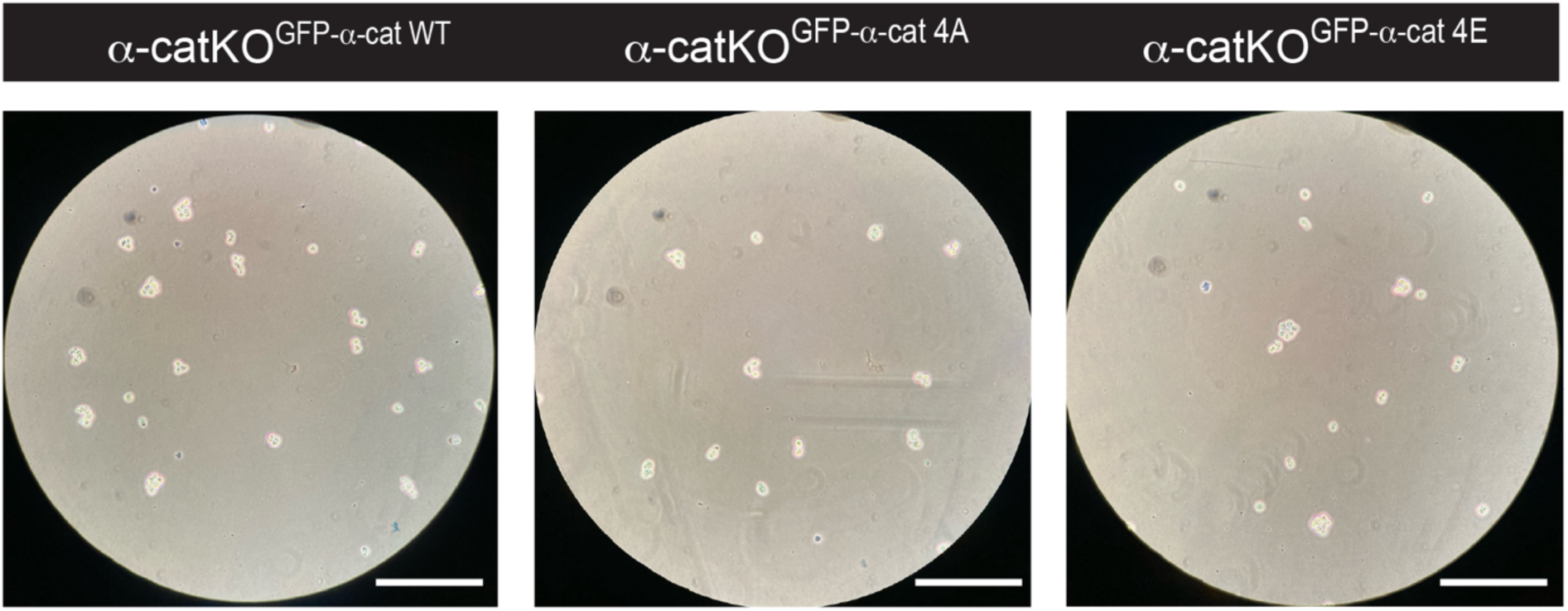
α-cat WT and phospho-mutant cell sizes are not intrinsically different. *[Related to Fig. 3]* Brightfield images taken on Nikon Eclipse TS100 with phone camera of α-cat mutant MDCK cells after trypsinization. Scale bar = 0.25mm.

**Video 1: Phospho-mimic α-cat restrains mitotic rounding.**

*[Related to Fig. 3]* Confocal images taken on AXR Nikon microscope of MDCK monolayer fixed and immuno-stained with antibodies to α-cat (green). DNA stained with Hoechst (blue). 4D image visualization on Imaris AI Microscopy Image Analysis Software was threshold, gated for voxels, and surface detail set between 0.2-0.5. Overlay 4D image analysis shows apical extension of nucleus during mitosis. Scale bar = 5μm.

**Fig. S3:**
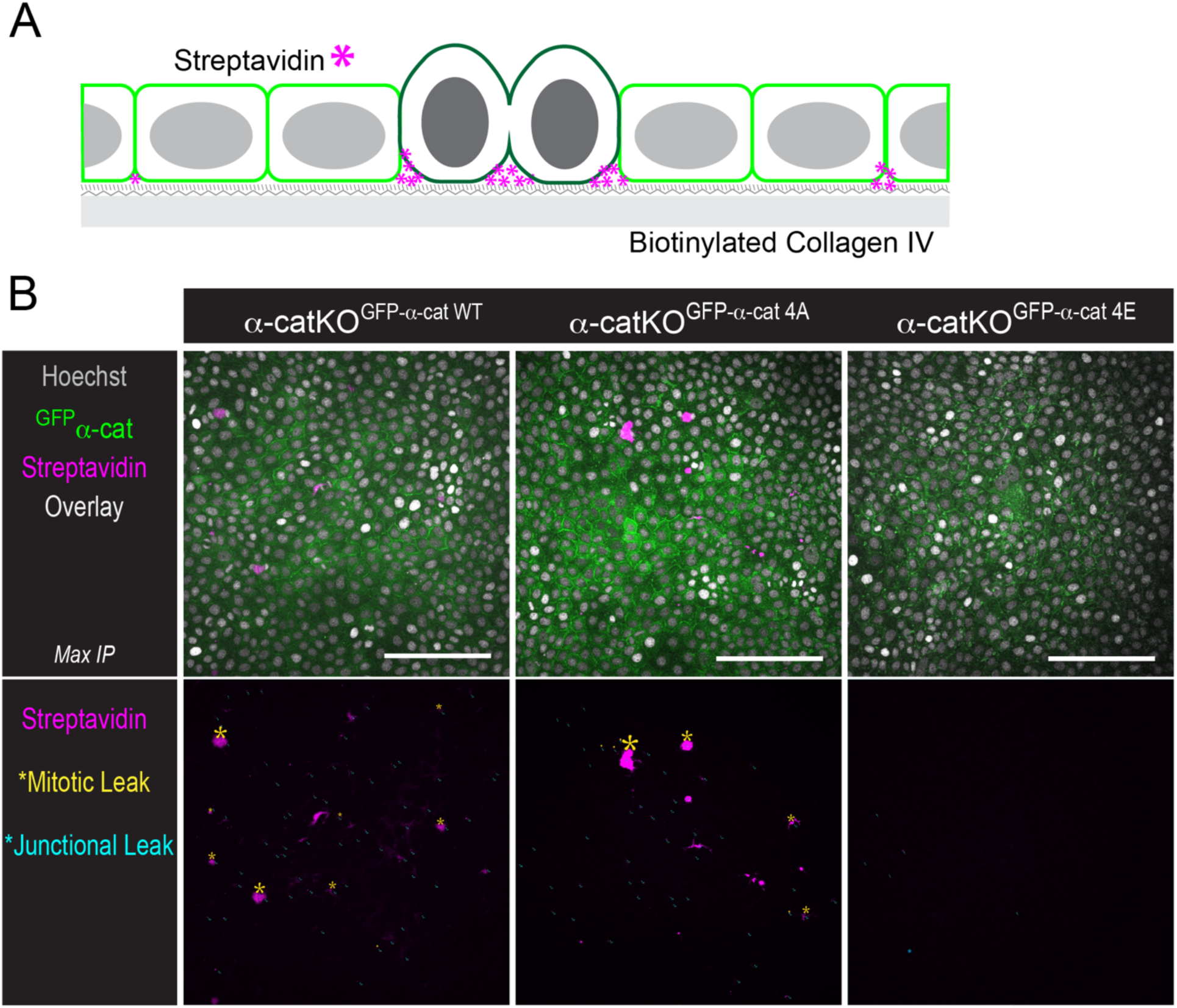
Phospho-mimic α-cat reduces barrier leak during mitotic rounding. *[Related to Fig. 4]* **(A)** Schematic of biotin-streptavidin permeability assay, where streptavidin (magenta asterisk) binds to biotinylated Collagen IV at barrier leaks during mitosis rounding. **(B)** Confocal image (z-stack maximum intensity projection of basal region) taken on Nikon AXR microscope of MDCK permeability assay fixed and immuno-stained for α-cat (native GFP, green), Hoechst (gray), and streptavidin (magenta). Hand tracing of mitotic leaks (yellow asterisk) and junctional leaks (tiny blue asterisks) showed reduced barrier leak in α-cat-KO^GFP-α-cat^ ^4E^ relative to α-cat-KO^GFP-α-cat^ ^WT^ or α-cat-KO^GFP-α-cat^ ^4A^ nascent monolayers. Scale bar = 100μm.

